# Reduction of a heme cofactor initiates N-nitroglycine degradation by NnlA

**DOI:** 10.1101/2022.06.19.496723

**Authors:** Kara A. Strickland, Ashley A. Holland, Alan Trudeau, Ilana Szlamkowicz, Melanie J. Beazley, Vasileios A. Anagnostopoulos, David E. Graham, Jonathan D. Caranto

## Abstract

The NnlA enzyme from *Variovorax sp.* strain JS1663 degrades the linear nitramine *N*-nitroglycine (NNG)—a natural product produced by some bacteria—to glyoxylate and nitrite (NO_2_^−^). Ammonium (NH_4_^+^) was predicted as the third product of this reaction. A source of non-heme Fe^II^ was shown to be required for initiation of NnlA activity. However, it was unclear if this Fe^II^ was being used as a metallocofactor or a reductant. This study reveals that NnlA contains a *b*-type heme cofactor. Reduction of this heme is required to initiate NnlA activity. Reduction can occur either by addition of a non-heme Fe^II^ source or by reduction with dithionite. Therefore, Fe^II^ is not an essential substrate for holoenzyme activity. Data are presented showing that reduced NnlA (Fe^II^-NnlA) can catalyze at least 100 turnovers. In addition, this catalysis occurred in the absence of O_2_. Finally, NH_4_^+^ was verified as the third product, accounting for the complete nitrogen mass balance. Size exclusion chromatography showed that NnlA is a dimer in solution. Additionally, Fe^II^-NnlA is oxidized by O_2_ and NO_2_^−^ and binds carbon monoxide (CO) and nitric oxide (NO). These are characteristics shared with PAS domains; NnlA was previously shown to exhibit homology with such domains. Providing further evidence, a structural homology model of NnlA was generated based on the structure of the PAS domain from *Pseudomonas aeruginosa* Aer2. The structural homology model suggested His^73^ is the axial ligand of the NnlA heme. Site-directed mutagenesis of His^73^ to alanine decreased the heme occupancy of NnlA and eliminated NNG activity, providing evidence that the homology model is valid. We conclude that NnlA forms a homodimeric heme-binding PAS domain protein that requires reduction for initiation of the activity.

**Importance:** Linear nitramines are potential carcinogens. These compounds result from environmental degradation of high-energy cyclic nitramines and as by-products of carbon capture technologies. Mechanistic understanding of the biodegradation of linear nitramines is critical to inform approaches for their remediation. The best understood biodegradation of a linear nitramine is NNG degradation by NnlA from *Variovorax sp.* strain JS 1663; however, it is unclear why non-heme iron was required to initiate enzymatic turnover. This study shows that non-heme iron is unnecessary. Instead, our study reveals that NnlA contains a heme cofactor, the reduction of which is critical for activating NNG degradation activity. These studies constrain the proposals for NnlA reaction mechanisms, thereby informing mechanistic studies of degradation of anthropogenic nitramine contaminants. In addition, these results will future work to design biocatalysts to degrade these nitramine contaminants.

## Introduction

Cyclic and linear (or aliphatic) nitramines are contaminants in soil and groundwater. Cyclic nitramines, such as hexahydro-1,3,5-trinitro-1,3,5-triazine, the high-energy compound RDX, are components of military-grade explosives. In addition to being products of RDX degradation, linear nitramines are by-products of some carbon capture technologies, formed when amines react with NO_x_ in the gas stream (1, 2).

RDX is a persistent soil contaminant at several explosives training facilities and manufacturing sites (3–5). Acute exposure to nitramines results in violent convulsions (6). Furthermore, the United States EPA lists RDX as an emerging contaminant and possible carcinogen. For these reasons, degradation pathways of RDX and other cyclic nitramines are well studied (7–13). One such pathway is initiated by a cytochrome P450 homolog, originally isolated from *Rhodococcus rhodochrous* strain 11Y, called XplA (14, 15). This enzyme reductively degrades RDX (**Fig. 1A**) (15–18). Under anaerobic conditions, the products are formaldehyde, nitrite (NO_2_^−^), and the one-carbon nitramine methylenedinitramine (MEDINA). Under aerobic conditions, RDX degradation produces the linear nitramine 4-nitro-2,4-diazabutanal (NDAB) instead. A transgenic *Arabidopsis* strain expressing the RDX degradation enzyme, XplA, has been engineered as a promising soil bioremediation strategy for RDX (3, 19).

**Fig. 1.**
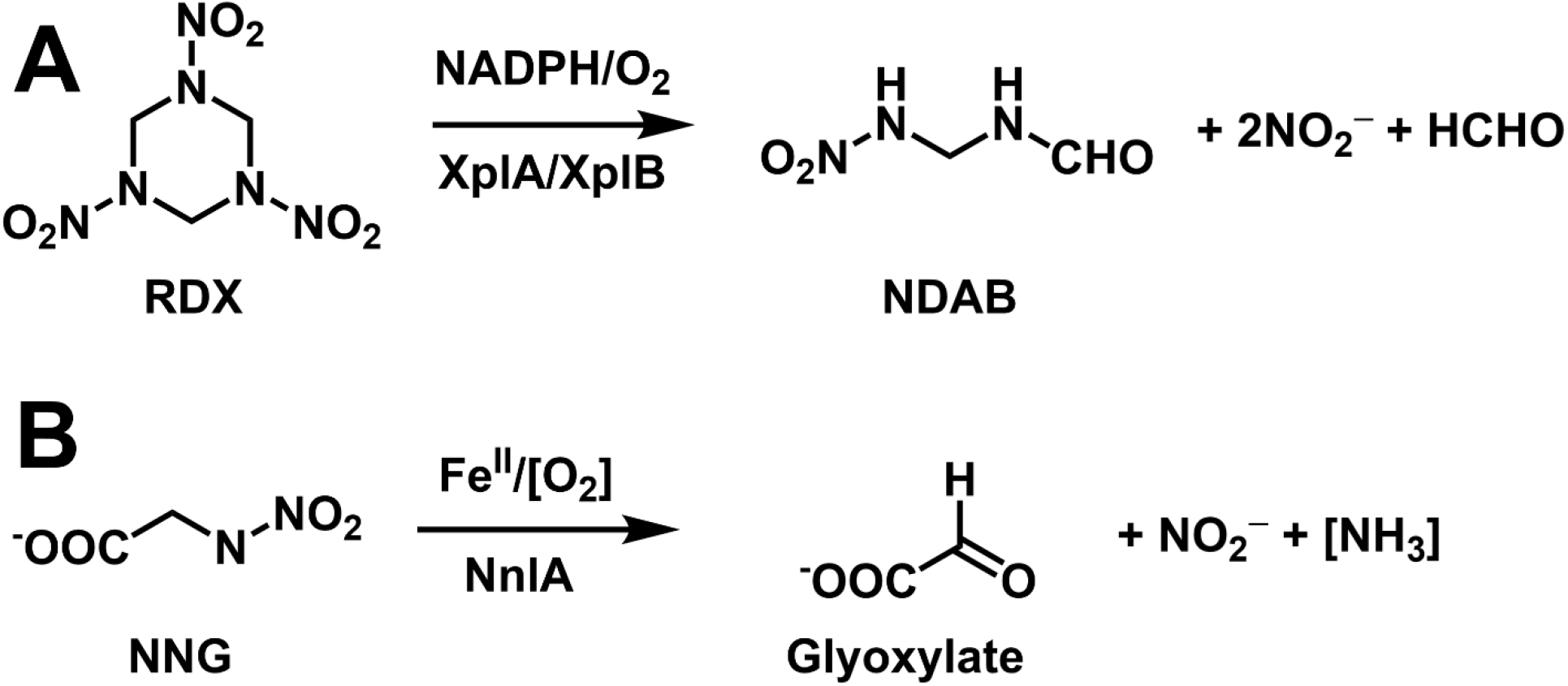
Product distributions of nitramine degrading enzymes, including A) aerobic degradation of RDX by XplA and B) degradation of NNG by NnlA. Reactants and products not yet verified are shown in square brackets.

Linear nitramines exhibit less acute toxicity than cyclic nitramines, but they have been shown to be skin and eye irritants (20, 21). Of greater concern, they are potential carcinogens (22–26). Linear nitramines often occur with their nitrosamine analogs, many of which are well-studied and potent carcinogens. Nitramines appear to be metabolized in a similar fashion as nitrosamines; specifically, their metabolism is initiated by hydroxylation by cytochromes P450 (22, 27–33). These hydroxylated products then degrade to form formaldehyde or alkylating agents, which are the proposed mutagenic agents. Linear nitramines are photostable (34). Therefore, biodegradation appears to be their main environmental degradation pathway. Understanding the mechanisms of linear nitramine biodegradation is necessary to inform effective bioremediation strategies and design new biocatalysts for this purpose.

Compared to cyclic nitramines, there is far less known regarding the biodegradation pathways and mechanisms of linear nitramines. Some linear nitramines produced by carbon capture, particularly those with hydroxyl functionalities, were shown to be biodegraded by bacterial samples from soil and water (35). The products of these degradations were not reported. Biodegradations of NDAB by the fungus *Phanerochaete chrysosporium* and the bacterium *Methylobacterium* sp. strain JS178 have been reported (36, 37). In both cases nitrous oxide (N_2_O) was produced. Initiation of the *P. chrysosporium* degradation was attributed to a manganese peroxidase. The mechanism of this degradation is unclear, but accumulation of N_2_O suggests it is initiated by cleavage between the 4N-3C bond of NDAB (**Fig. 1A**)

The best characterized linear nitramine biodegradation pathway is an enzymatic pathway that degrades the natural product *N*-nitroglycine (NNG). This linear nitramine is produced by some bacteria including *Streptomyces noursei* (38, 39). The physiological function of NNG is unknown, but it exhibits toxicity towards Gram-negative bacteria and plants (38, 40). In addition, *in vitro* experiments have shown that NNG inhibits the ubiquitous succinate dehydrogenase enzyme (41). *Variovorax* sp. strain JS 1663 was enriched by its ability to use NNG as a sole carbon and nitrogen source (42). An NNG lyase (NnlA) was discovered to be essential for this phenotype. *In vitro* assays of NnlA showed it degraded NNG, producing glyoxylate and NO_2_^−^ (**Fig. 1B**). A second nitrogenous product remains unidentified but was predicted to be ammonium (NH_4_^+^). Previous characterization of NnlA showed no evidence for a redox cofactor, such as a heme or flavin.(43, 44) Therefore, NH_4_^+^ as the final product was consistent with a redox-neutral degradation pathway and the apparent lack of a redox cofactor requirement. However, NnlA was shown to be homologous to PAS domain proteins, which typically bind heme or flavin. Furthermore, initiation of activity for heterologously expressed, purified NnlA required addition of ferrous ammonium sulfate ([Fe(NH_4_)_2_(SO_4_)_2_]). This Fe^II^ may have reconstituted a non-heme iron-containing active site capable of activating dioxygen (O_2_)—a well-known role for non-heme iron sites(45). An iron-dependent redox-mediated NNG degradation pathway by NnlA could be envisioned. To test this hypothesis, the role of Fe^II^ for NnlA activity needs to be clarified.

The purpose of this study was to identify the role of Fe^II^ in NnlA activity. En route to solving this question, we found that NnlA binds a heme cofactor. As with non-heme iron, heme cofactors are well-established to enable O_2_ activation (46). Alternatively, heme may enable a reductive degradation pathway as observed for RDX degradation by XplA. We investigated the role of the NnlA heme cofactor, fully characterized the nitrogen mass balance of NNG degradation by NnlA, and the O_2_-dependence of turnover. The cumulative data herein explains why Fe^II^ was needed to initiate activity in the prior work and shows it is not essential for activity. Finally, we provide evidence that NnlA shares several characteristics with heme-binding PAS domain proteins. The results provide insight into the mechanism of linear nitramine degradation and will aid in developing remediation strategies and engineering new enzymes for bioprocessing.

## Material and Methods

### General reagents and protocols

Isopropyl β-D-1-thiogalactopyranoside (IPTG) and 5-aminolevulinic acid (5-ALA) were purchased from GoldBio. NNG was purchased from AAblocks. General buffers and media components were purchased from Fisher Scientific or VWR. Stock dithionite concentrations were determined by UV-visible absorbance at 318 nm (ε_318_ = 8000 M^-1^cm^-1^). The nitric oxide generator PROLI-NONOate was purchased from Cayman Chemicals. Stocks of PROLI-NONOate were prepared by dissolving approximately 10 mg of PROLI-NONOate in 10 mM NaOH and quantified by measuring the absorbance at 250 nm (ε_250_ = 6500 M^-1^cm^-1^) or 252 nm (ε = 8400 M^-1^cm^-1^), respectively. NO gas was purified prior to use by bubbling into degassed 10 mM NaOH in a septum-sealed container. Buffers were degassed in septum-sealed glass bottles by 3x vacuum/N_2_ gas purge cycles on a Schlenk line connected using a 22 G needle punctured through the septum. Water used for all solutions was of 18.2 MΩ·cm resistivity from a Barnstead Nanopure (Thermo Fisher Scientific). Solvents for LC-MS experiments were of at least HPLC grade and contained 0.1% *v*/*v* formic acid. Recombinant TEV protease was expressed and purified as previously described (47).

### Mutagenesis of nnlA

Site directed mutagenesis of the *nnlA* gene was performed to produce the variant protein NnlA H73A. Primers were designed for an “Round-the-horn” mutagenesis (**Table S1**) (48). The standard protocol was performed using Phusion polymerase and an annealing temperature of 63°C. The ligated reaction mixture was transformed into competent *E. coli* DH5α cells and plated on terrific broth (TB) agar plate containing 0.1 g/mL ampicillin. Colonies were selected and used to inoculate 5-mL Luria Broth (LB) cultures with 0.1 g/mL ampicillin, which were grown overnight at 37 °C. DNA was extracted from these cultures and analyzed by DNA sequencing (GeneWiz) to verify the mutation.

### Expression and purification of NnlA and variant

Gene expression and affinity purification of NnlA protein using the pDG708 expression vector was performed as previously described (42). Alternatively, NnlA could be expressed from T7 Express cells transformed with pDG750, which contains the *nnlA* gene with an N-terminal His_10_-tag, which is codon-optimized for expression in *E. coli*. NnlA with high heme occupancy was expressed by growth of these transformants in 4 x 1-L flasks of TB with 0.1 g/L of ampicillin at 37 °C. At an OD_600_ of 2, the temperature was decreased to 20 °C and NnlA expression induced with 100 mg/L of isopropyl ß-D-1-thiogalactopyranoside (IPTG), 1 g/L of ferric ammonium citrate (FAC) and 0.8 g/L 5-aminolevulinic acid (5-ALA). The cultures were grown for another 24 hours. Cells were pelleted by centrifuge at 6353 *g* yielding 17 g/L of wet cell mass. The pellet was either lysed immediately or stored frozen at −60 °C.

Cells were lysed by resuspending the cell pellet in a 1:2 (*m*/*v*) ratio of cells to Ni-buffer A (100 mM Tris-HCl, 100 mM NaCl at pH 7.6). The cell suspension was sonicated at 20 % amplitude for 10 minutes (10 seconds pulse, 10 seconds pause) for 3 cycles on ice. The cell debris was pelleted in a centrifuge at 53,000 *g* yielding a clear cherry-brown lysate. His_10_-NnlA was loaded on a Ni-NTA HTC column (GoldBio), washed with 2 column volumes of Ni-buffer A, and eluted using a 6 column-volume gradient of 0 to 100% Ni-buffer B (100 mM Tris-HCl, 100 mM NaCl, 1 M imidazole at pH 7.6). Reddish-brown fractions were evaluated for relative heme incorporation using the ratio of 412 nm to 280 nm based on UV-visible absorption spectra. We found that fractions with a ratio A_412_/A_280_ > 1.0 had the highest heme occupancies. Therefore, these fractions were evaluated for purity using SDS-PAGE. Pooled fractions were buffer exchanged into Ni-buffer A. Concentrated protein was aliquoted into microcentrifuge tubes and stored at −60 °C. NnlA H73A was expressed and purified as described above.

### NnlA characterization and spectroscopy

NnlA was quantified using a bicinchoninic acid assay (Thermo Scientific). Total heme and non-heme iron was quantified using a literature iron assay that allows for release and subsequent detection of heme-ligated iron (49). Non-heme iron was specifically quantified using a ferrozine assay (50). Differentiation of the bound heme type was determined using the pyridine hemochromagen assay (51). Dissolved metal concentrations were determined using a Thermo Fisher Scientific iCAP-Qc inductively coupled plasma mass spectrometer (ICP-MS) with QCell technology and operated in kinetic energy discrimination (KED) mode of analysis with helium as the collision gas. Calibration, internal, and quality control standards (Inorganic Ventures) were prepared in 2% (v/v) HNO_3_ and calibration standards were prepared at concentrations of 1 to 1000 µg L^-1^ (ppb).

To characterize the oligomer state of NnlA. The His_10_-tag was cleaved from NnlA using recombinant TEV protease as previously described (47). The His_10_-NnlA was incubated with TEV protease at 4°C for 72 hours. The digested protein was passed over a 2-mL Ni-HTC (GoldBio) gravity column to separate the tag-free NnlA, remaining His_10_-NnlA, His_7_-TEV, and cleaved His_10_-tag. The brown flowthrough containing tag-free NnlA was concentrated, and 500 μL was then passed over a Superdex TM200 10/300 GL analytical gel filtration column. The column was preequilibrated with 100mM Tris-HCl, 100 mM NaCl at pH 7.8 at a flow rate of 0.1 mL/min. A standard mass curve was generated by passing a premade BioRad Gel filtration standard (#1511901) and blue dextran.

UV-visible absorption spectra of NnlA oxidation states and gas-bound forms were collected in an anaerobic glovebox (Vacuum Atmospheres Co.) atmosphere using an Ocean Optics USB2000+ UV-visible absorption spectrometer. Samples containing 10 μM NnlA in degassed buffer were titrated with stock sodium dithionite until the 430 nm absorbance of Fe^II^ NnlA no longer increased. The ferrous-carbon monoxide and ferrous-nitrosyl NnlA adducts were generated by addition of either CO gas or purified NO gas to an anaerobic sample of 10 μM Fe^II^ NnlA contained in a septum sealed quartz cuvette. Spectra of Fe^II^ NnlA reacted with NNG or NO_2_^−^ were collected after reaction of 2.7 μM Fe^II^ NnlA with 133 μM NNG or nitrite. Single wavelength traces were collected by monitoring 415, 417, or 437 nm absorbance every 4.8 seconds for the duration of the reaction.

### General LC-MS methods

LC-MS analysis was performed using an Agilent 1260 LC stack equipped with a Zorbax RX-C18 column (5 μm, 4.6 x 150 mm) and connected to an Agilent 6230 TOF mass spectrometer with electrospray ionization (ESI). Analyses used an isocratic mixture containing 65% water, 25% acetonitrile, and 10% isopropanol at a flow rate of 0.5 mL/min. The mass spectrometer was run in negative ion mode with a probe voltage of 4500 V and fragmentation voltage of 175 V. To monitor NNG and glyoxylate, extracted ion chromatograms were obtained at *m*/*z* 119.0 and *m*/*z* 73.0, respectively. Concentrations of NNG and glyoxylate in samples were determined using calibration curves of standards containing 65, 125, 250, 500, and 750 µM NNG or glyoxylate in 10 mM tricine buffer.

### Nitrogen assays

Ammonium concentrations were determined using a glutamate dehydrogenase assay (Sigma-Aldrich) kit using the manufacturer’s instructions. Nitrite concentrations were determined by reacting 100 μL aliquots of reaction sample with 50 μL of deoxygenated Griess reagent R1 (Cayman Chemical, 1% sulfanilamide in 5% H_3_PO_4_) followed by addition of 50 μL of deoxygenated Griess reagent R2 (Cayman Chemical, 0.1% napthylethylenediamine dihydrochloride in water). The NO_2_^−^ concentration was determined by using the molar absorption coefficient of the Griess-treated sample (ε_542_ = 50 mM^-1^cm^-1^) or from a standard curve generated from reactions with known concentrations of NO_2_^−^.

Samples were analyzed by ion chromatography for nitrite and nitrate using a Dionex Integrion High-Pressure Ion Chromatography (ThermoScientific) equipped with a 4 mm anionic exchange column (IonPac AS20), suppressor (Dionex ADRS 600 Suppressor) and a conductivity detector, operated at constant voltage (4.0V). The sample loop was 20 μL and was first degassed in an internal oven at 30 °C and then carried through the column by 35 mM NaOH (ultrapure, carbonate free, Acros Organics). The elution times under the conditions studied were 4 min and 4.9 min for nitrites and nitrates, respectively.

### Preparation of NNG decomposition stoichiometry samples

Stoichiometry samples were prepared in an anaerobic glovebox using degassed buffers. Samples containing 10 µM as-isolated (NnlA_as-iso_) or Fe^II^ NnlA were mixed with 500 µM NNG as needed for each sample. The dithionite control was prepared by mixing 500 µM NNG with 10 µM Na_2_S_2_O_4_. Reduced samples contained 10 µM Na_2_S_2_O_4_, 10 µM NnlA_as-iso_, and 500 µM NNG in 10 mM tricine buffer, pH 8. Samples reacted with O_2_ were prepared in two parts: 250 μL of 1 mM NNG remained outside of the glovebox under oxic conditions, while 250 μL of reduced protein was prepared inside the anaerobic glovebox by mixing NnlA and Na_2_S_2_O_4_ to a final concentration of 20 μM each. The 250 μL aliquot of protein was removed from the box and added quickly to the 250 μL aliquot of oxygenated NNG in buffer with a final concentration of 20 mM potassium phosphate and pH 7.2. Every sample prepared was incubated for 30 minutes regardless of conditions.

## Results

### Characterization of NnlA and its heme cofactor

Recombinant NnlA with an N-terminal protease-cleavable decahistidine tag (His_10_-NnlA) was produced in *E. coli* T7 Express cells. The His_10_-NnlA protein was purified from the lysate by a Ni-NTA column. The theoretical mass of His_10_-NnlA was calculated as 20,526 Da. While a monomer band was observed in the SDS-PAGE of protein purified by a Ni NTA column, several other higher molecular weight bands also appeared (**Fig. S1A**). These higher molecular weight bands have weights consistent with NnlA oligomers.

To determine the solution oligomerization state of NnlA, TEV protease-cleaved NnlA was analyzed using size exclusion chromatography in 10 mM Tris-HCl at pH 7.6 (**Fig. S1B**). The chromatogram of TEV-cleaved NnlA exhibited two peaks. The first peak elutes in the void volume suggesting a higher oligomer with apparent molecular mass of 200 kDa or greater. The second peak corresponds to a molecular mass of 44 kDa, which is consistent with an NnlA homodimer. There was no appearance of a monomer peak. The results suggested that NnlA is a homodimer in solution, with some of the NnlA appearing as a larger aggregate.

The UV-visible absorption spectrum of as isolated His_10_-NnlA exhibited a feature centered at 412 nm consistent with the presence of heme (**Fig. 2**). Supplementing expression cultures with 5-aminolevulinic acid (5-ALA), a heme precursor, resulted in preparations with a 6-fold higher molar absorption coefficient at 412 nm. These results indicate that addition of 5-ALA increases the occupancy of heme in recombinant NnlA. These preparations routinely contained 0.87 ± 0.21 total Fe atoms per NnlA monomer (**Table S2**). By contrast, negligible non-heme iron was detected by ferrozine assay. UV-visible absorption of samples of NnlA and myoglobin treated with the pyridine hemochromagen assay were nearly identical (**Fig. S2**). These results are consistent with the binding of a *b*-type heme. Other transition metals were detected in His_10_-NnlA samples by ICP-MS (**Table S3**). However, most of these were sub-stoichiometric compared to the protein and are not expected to bind specifically to the protein. Nickel concentrations were high and likely due to the use of the Ni-NTA column during purification (52). As discussed above, amino acid sequence alignment shows that NnlA contains a PAS domain. The dimerization of the monomer and binding of a heme cofactor are common traits of these domains (43, 44, 53).

**Fig. 2.**
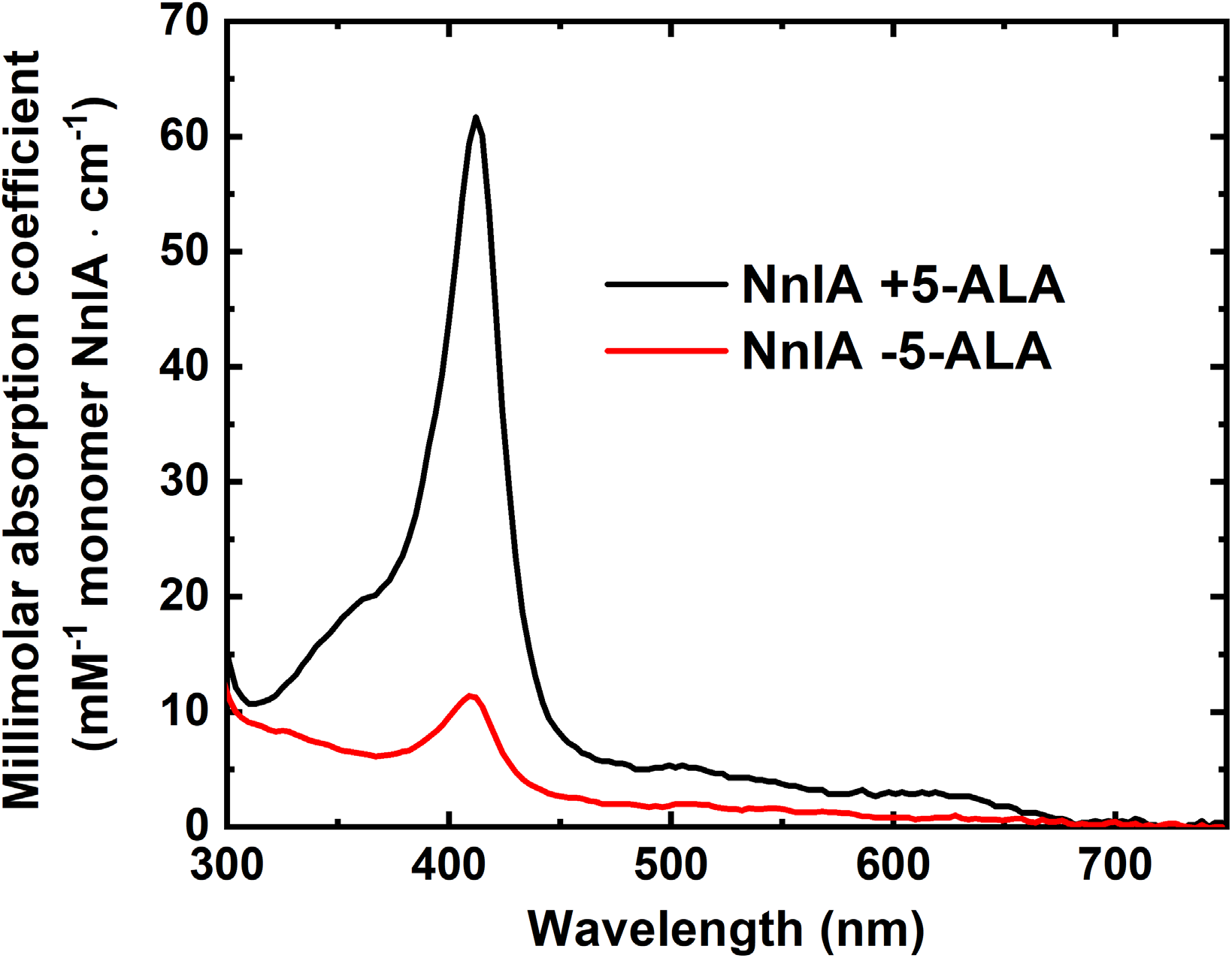
UV-visible absorption spectra of as isolated NnlA expressed in the absence (red trace) or presence (black trace) of 5-ALA. Both spectra were measured in 50 mM phosphate buffer at pH 7.2.

### Reduced NnlA reacts with O_2_, NO, and CO

The role of the heme cofactor in PAS domains is often a sensor of redox environment or of gas molecules such dioxygen (O_2_), or nitric oxide (NO) (53–55). Therefore, we characterized the reactions of paramagnetic gas molecules with reduced NnlA. Reduction of the NnlA heme was monitored by UV-visible absorption spectrophotometry (**Fig. 3**). The as isolated His_10_-NnlA exhibited a Soret feature at 413 nm with broad absorbance in the 500–700 nm region including poorly resolved peaks near 502 and 636 nm. The absorption spectrum is reminiscent of Fe^III^ hemoglobin or myoglobin (56). Therefore, the as isolated His_10_-NnlA is hereafter termed Fe^III^-NnlA. Reduction of the sample was achieved by titration of Fe^III^-NnlA with dithionite until the absorbance at 413 nm no longer decreased. Alternatively, the protein could be reduced by addition of an excess of Fe(NH_4_)_2_(SO_4_)_2_ (**Fig. S3**). Under either reduction condition, the resulting reduced sample exhibited a Soret band at 437 nm and formation of a Q-band maximum at 560 nm. This spectrum of reduced NnlA resembles that of Fe^II^ myoglobin (56). Therefore, these features are hereafter attributed to Fe^II^-NnlA and are achieved using dithionite as the reductant.

**Fig. 3.**
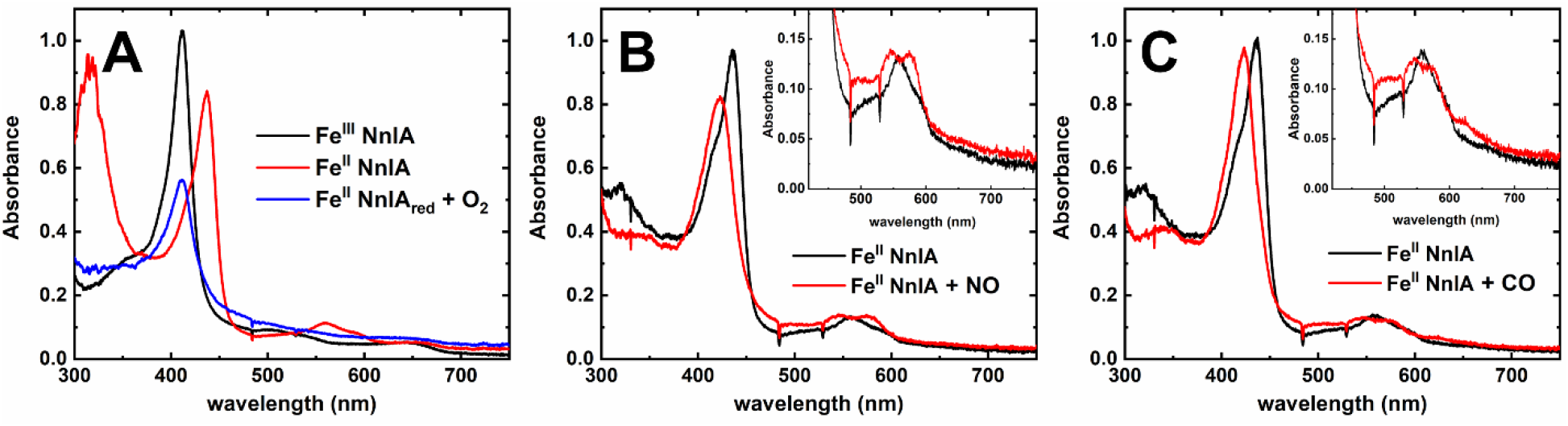
Treatment of Fe^II^-NnlA with A) O_2_, B) NO or C) CO. Conditions before addition of gas are 10 μM Fe^II^-NnlA in either in 100 mM Tris-HCl pH 7.6 (panel A) or 100 mM tricine buffer at pH 8 (panels B and C).

The reaction of O_2_, CO, or NO with Fe^II^-NnlA was monitored by UV-visible absorption spectrophotometry. There were no noticeable changes in the UV-visible absorption spectra when O_2_, CO, or NO were added to Fe^III^-NnlA (**data not shown**). By contrast, each of these gases readily reacted with Fe^II^-NnlA (**Fig. 3**). Exposure of 10 μM Fe^II^-NnlA to air resulted in the loss of Fe^II^-NnlA absorption features with a concomitant rise in Fe^III^-NnlA absorption features over several minutes. The intensities of these features were decreased compared to those observed for the protein prior to reduction (**Fig. 3A**), suggesting that the observed oxidation of Fe^II^-NnlA by O_2_ destroys some of the heme. Optical features consistent with the Fe^II^-O_2_ were not observed, but it is expected to be an intermediate en route to heme oxidation.

Addition of NO gas to a deoxygenated sample of Fe^II^-NnlA resulted in immediate appearance of new absorbance features at 421, 550, and 580 nm. This spectrum is consistent with that for the ferrous-nitrosyl ({FeNO}^7^ by Enemark-Feltham notation (57)) adduct of myoglobin (**Fig. 3B**) (56). Meanwhile, addition of CO to Fe^II^-NnlA causes appearance of new absorption features at 425, 542, and 568 nm. This spectrum is similar to that of the CO adduct of myoglobin (58), thereby indicating that Fe^II^-NnlA binds CO (**Fig. 3C**).

### NnlA activity requires heme reduction and does not require O_2_

The NNG degradation activity of heme-occupied Fe^III^-NnlA was first tested. Samples containing Fe^III^-NnlA and NNG were analyzed with LC-ESI-MS to monitor for decomposition of NNG (*m*/*z* 119.0) and formation of glyoxylate (*m*/*z* 73.0). Extracted ion chromatograms (EICs) monitoring these molecular anions are shown in **Fig. 4**. These Fe^III^-NnlA samples showed no evidence for NNG decomposition to form glyoxylate, suggesting Fe^III^-NnlA needs to be activated to exhibit activity. As shown above, Fe(NH_4_)_2_(SO_4_)_2_—required to initiate activity in prior work—was able to reduce the NnlA heme. Therefore, we posited that NnlA activity is dependent on reduction of the heme instead of the presence of a ferrous iron source.

**Fig. 4.**
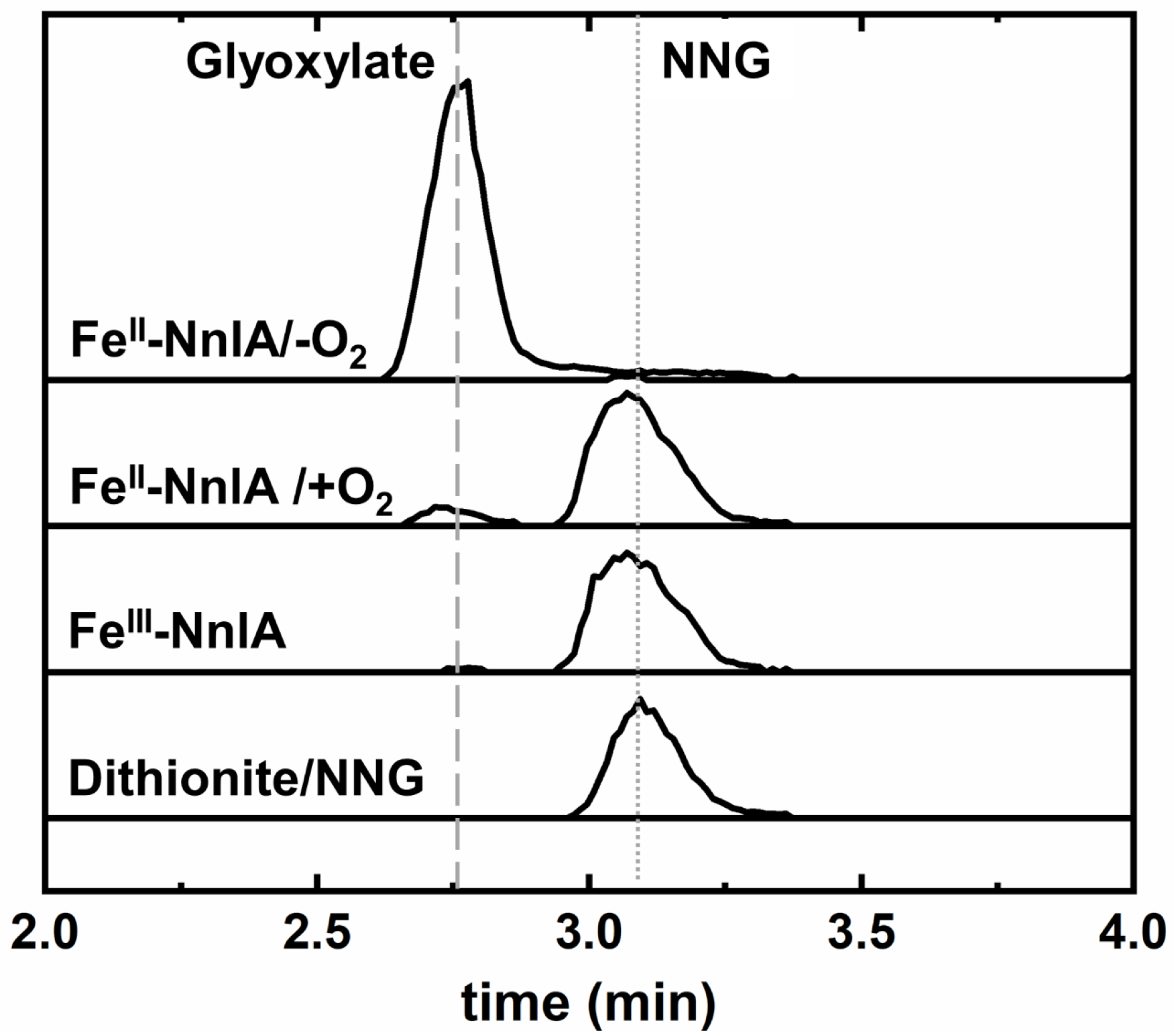
Overlaid representative LC-MS EICs monitoring molecular anions of NNG (*m*/*z* 119.0) and glyoxylate (*m*/*z* 73.0) in samples containing 500 μM NNG and 10 μM dithionite (Dithionite/NNG), 10 μM as isolated NnlA (Fe^III^-NnlA), or 10 μM reduced NnlA (Fe^II^-NnlA). Samples were incubated for 30 minutes at room temperature in deoxygenated 20 mM phosphate buffer, pH 7.2. The Fe^II^-NnlA/O_2_ sample was incubated in air-saturated buffer. Dashed and dotted gray lines indicate elution time of glyoxylate and NNG in standard solutions.

To test this hypothesis, Fe^II^-NnlA was incubated with NNG. These samples were prepared in an anaerobic glovebox test the need for O_2_ for the reaction. Anaerobic samples containing 10 µM Fe^II^-NnlA and 500 µM NNG exhibited complete degradation of the NNG to form glyoxylate (**Fig. 4**). For comparison, samples containing only dithionite and NNG exhibited no degradation of NNG. This result showed that the NnlA heme needs to be in the Fe^II^ oxidation state to activate NNG decomposition activity. The need for reduction of the heme to initiate NnlA activity unambiguously showed that the heme cofactor is necessary for NnlA activity, and its appearance is not an artifact of recombinant expression. Furthermore, this activation could be achieved without the need for Fe(NH_4_)_2_(SO_4_)_2_.

These experiments also precluded the hypothesis that NNG degradation by NnlA proceeds by a reductive or oxidative pathway. First, 10 µM Fe^II^-NnlA performed 50 turnovers under these conditions. This catalytic NNG degradation shows that electron transfer from the heme to the NNG is not necessary, and thereby eliminated the possibility of a reductive NNG degradation pathway. In addition, complete NNG degradation was observed in the absence of O_2_. This result precluded the possibility that O_2_ activation by the NnlA heme is required for NNG degradation. In fact, simultaneous addition of O_2_ and NNG resulted in less NNG degradation, indicating O_2_ inhibits the reaction. These combined observations strongly suggested that NNG degradation by NnlA is redox-neutral.

### Determination of the nitrogen mass balance

To verify that NNG degradation by NnlA is redox neutral, we determined the mass balance of the NNG degradation. Samples containing 10 µM Fe^II^-NnlA and 1000 μM NNG in 10 mM tricine buffer, pH 8 were prepared under anaerobic conditions and incubated at 20 °C for 30 minutes. In parallel, NNG and glyoxylate were analyzed by LC-ESI-MS, NO_2_^−^ was quantified by Griess assay or ion chromatography, and NH_4_^+^ was assayed by a L-glutamate dehydrogenase coupled assay. The EICs showed that the NNG in these samples was completely degraded with concomitant appearance of 1210 ± 290 μM glyoxylate, accounting for 100% of the carbon mass balance. Parallel analysis of nitrogenous products showed the appearance of 810 ± 40 µM NO_2_^−^ and 1050 ± 100 µM NH_4_^+^ in these samples. Ion chromatography showed no evidence for the presence nitrate (**Table S4**), providing further evidence against NNG degradation proceeding by an oxidative pathway. The total nitrogen products accounts for 93% of the nitrogen mass balance. These results are consistent with the following reaction stoichiometry:

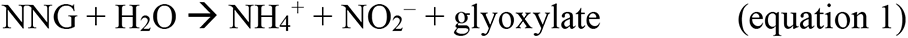

This reaction stoichiometry of NNG degradation by NnlA is redox neutral.

### NO_2_^−^ oxidizes Fe^II^ NnlA

Given that the reaction stoichiometry was consistent with a redox-neutral process, it was not expected that NNG would oxidize Fe^II^-NnlA. However, samples containing 3 μM Fe^II^-NnlA reacted with 133 μM NNG under anaerobic conditions resulted in oxidation of Fe^II^-NnlA within several hours as monitored by UV-visible absorption spectrophotometry (**Fig. 5A**). The final spectrum of this reaction exhibited spectral features Q-band features consistent with formation of the heme {FeNO}^7^ shown in **Fig. 3B**. This observation suggested NNG degradation resulted in formation of some NO, which subsequently bound to Fe^II^-NnlA to form the observed heme-nitrosyl adduct. Heme centers are well known to reduce nitrite to NO (59); therefore, we posited that the product NO_2_^−^ was responsible for oxidizing the heme center and not NNG.

**Fig. 5.**
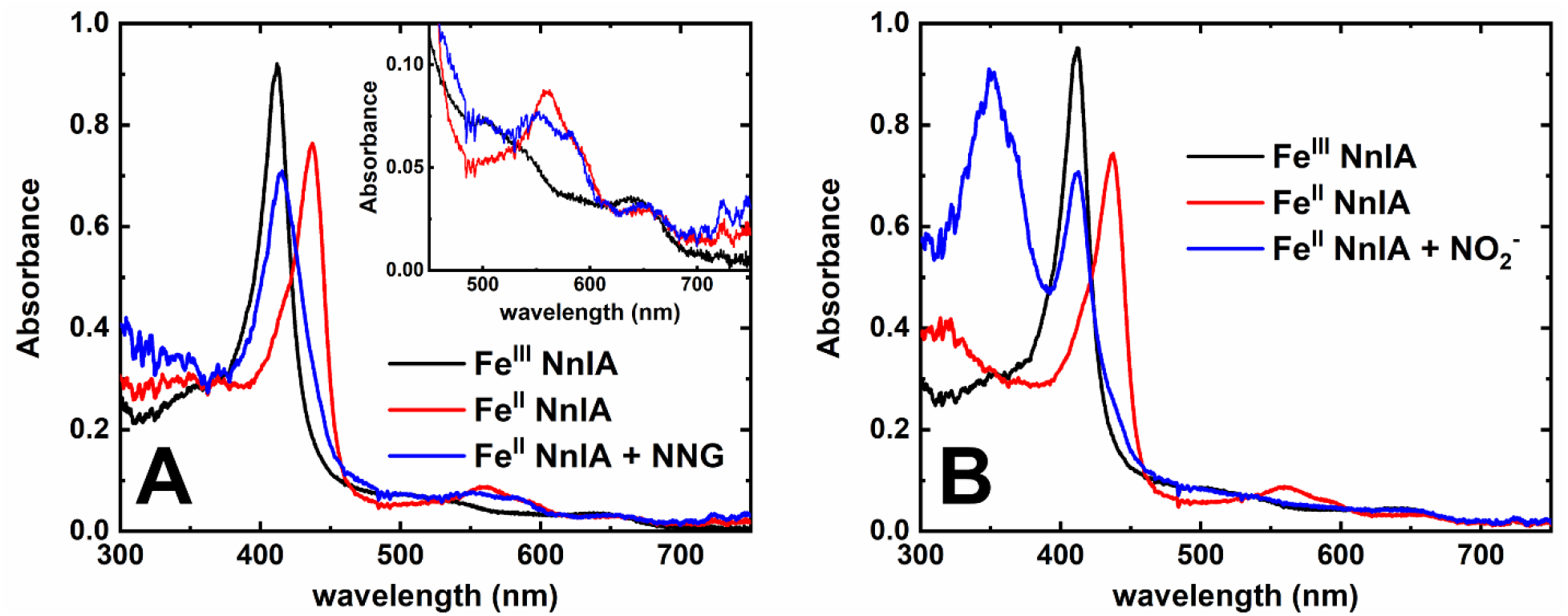
UV-visible spectra of reduced NnlA treated with either NNG (Panel A) or NO_2_^−^ (Panel B) under anaerobic conditions. Reaction conditions: (Panel A) 3 µM Fe^II^-NnlA, 133 µM NNG, in deoxygenated 20 mM phosphate, pH 7.5 incubated for 5 hours at room temperature; (Panel B) 3 μM Fe^II^-NnlA with 133 μM NO_2_^−^ in deoxygenated 20 mM phosphate, pH 7.5 at room temperature.

Indeed, addition of 133 µM NO_2_^−^ to 3 μM of Fe^II^-NnlA under anaerobic conditions oxidized the protein to Fe^III^-NnlA (**Fig. 5B**). We note that in this experiment, there is no evidence for formation of an {FeNO}^7^ species. Nevertheless, the time course of oxidation of the Fe^II^-NnlA center, monitored by the decrease in 437 nm, was nearly identical whether in the presence of NO_2_^−^ or NNG (**Fig. S4**). These combined results indicated that NO_2_^−^, a product of NNG degradation, oxidized Fe^II^ NnlA to Fe^III^ NnlA, thereby inactivating the protein. In other words, the data show that NnlA was product inhibited.

### Structural homology model of NnlA

To date, there are no published crystal structure models of NnlA. However, the accumulated evidence suggests that NnlA is a PAS domain protein. To provide structural prediction of NnlA, we generated a structural homology model using SWISS-MODEL (**Fig. 6**). The model was selected by first allowing SWISS-MODEL to self-select recommended templates. Models were then generated on all structural templates containing a heme cofactor, consistent with the experimentally observed heme in NnlA. This method resulted in the generation of 17 structural models. Out of the 17 models, 16 were based on previously characterized PAS domains including those from DOS, FixL, and Aer2 protein complexes. The best generated model threaded the NnlA sequence onto the crystal structure of the cyanide (CN)-bound PAS domain of *Pseudomonas aeruginosa* Aer2 (PDB ID: 3VOL) and had a GMQE score of 0.31. All other models exhibited a GMQE score between 0.03 and 0.18.

**Fig. 6.**
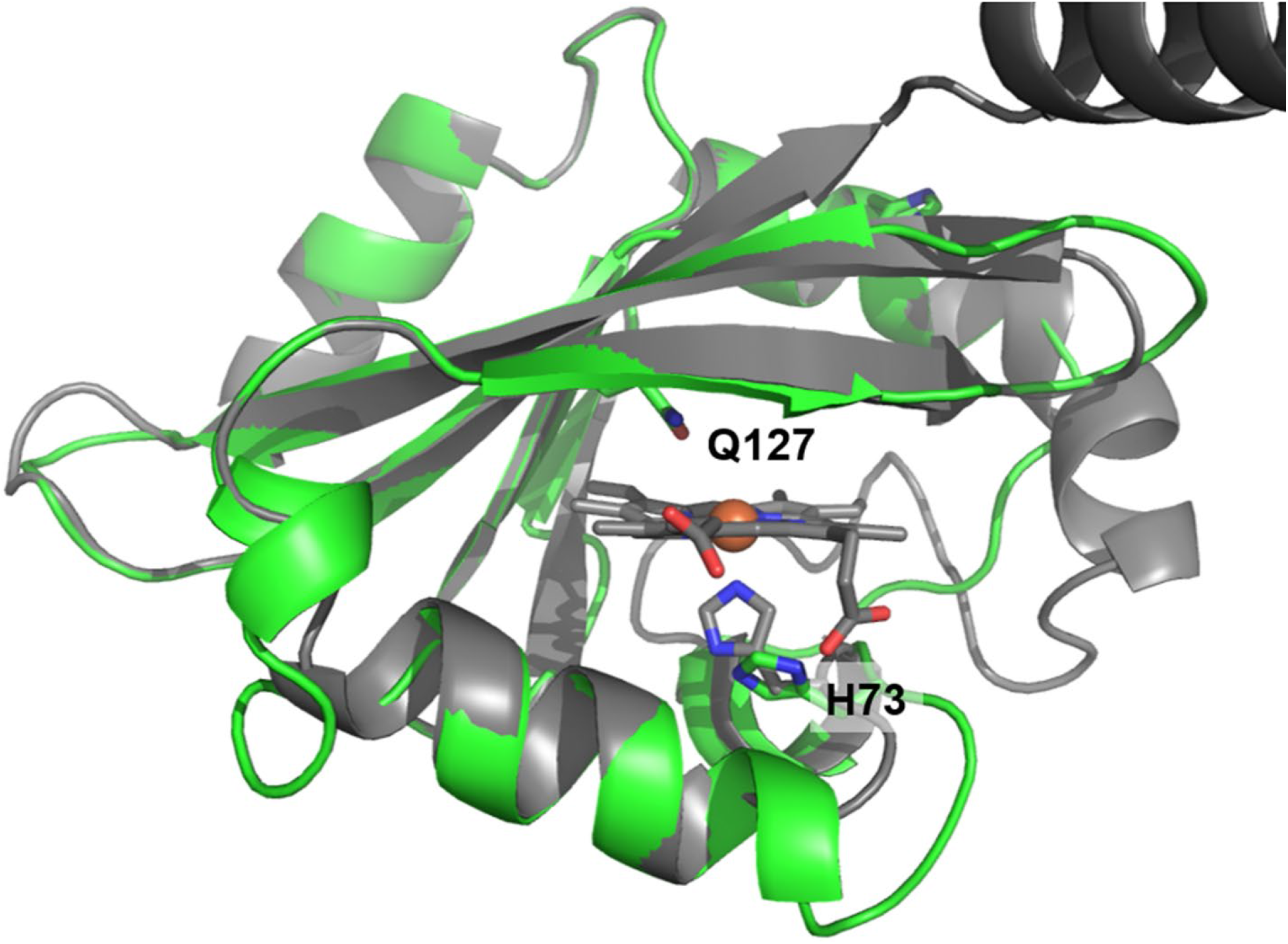
Structural homology model of NnlA overlayed on the crystal structure of the CN^−^ complexed PAS domain from *Pa* Aer2 (PDB: 3VOL; grey). Peptide backbone of NnlA structural homology model shown as green cartoon and *Pa* Aer2 backbone shown as grey cartoon. Iron atom shown as orange sphere, oxygen and nitrogen atoms shown in red and blue, respectively. Structural homology model was generated by SWISS-MODEL.

We tested the homology model by generating an NnlA variant with decreased heme occupancy. In *Pa* Aer2, the heme is coordinated by an axial histidine ligand (60). Overlaying the NnlA structural homology model on the *Pa* Aer2 structure reveals that H73 of NnlA is in a similar position as the axial histidine coordinated to the *Pa* Aer2 heme (**Fig. 6**). To validate the structural homology model and H73A NnlA variant was generated by site-directed mutagenesis. The variant protein recombinantly expressed and purified as described for wild-type NnlA. The total iron concentration per monomer protein (0.35 ± 0.08 μM Fe/monomer) of the H73A NnlA was less than half that for wild-type NnlA (**Table S2**). Furthermore, there is a large amount of non-heme iron in the H73A NnlA samples, suggesting that the total iron assay overestimates the heme occupancy. Regardless, the iron analyses suggest that the H73A mutation adversely affects the binding of the heme cofactor. This conclusion is supported by UV-visible absorption spectra, which showed that the purified H73A NnlA exhibited an 8-fold lower molar absorption coefficient at 410 nm that that of the wild-type NnlA (**Fig. 7**). Finally, the variant protein lacks NNG degradation activity. There is no difference in the amount of NNG or glyoxylate in samples of Fe^II^-H73A NnlA compared to Fe^III^-H73A NnlA samples incubated with NNG for 30 minutes (**Fig. S5**). The combined results validate the proposed heme binding site from the structural homology model of NnlA.

**Fig. 7.**
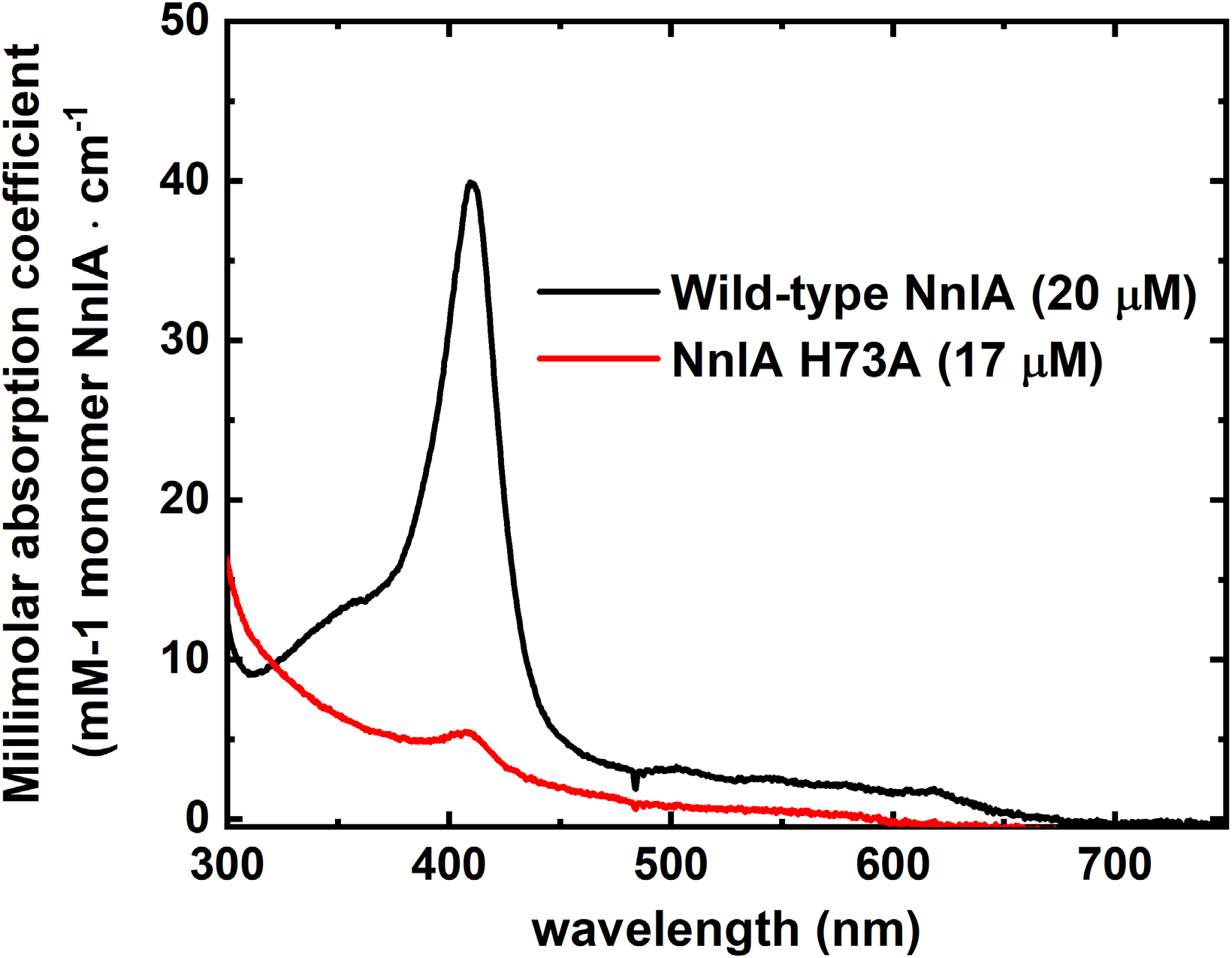
UV-visible absorption spectra of 20 μM wild-type or 17 μM H73A NnlA. All samples prepared in 100 mM tricine buffer, pH 8.

## Discussion

The conclusions of the data presented above are summarized in **Scheme 1**. A critical first conclusion is that NnlA contains a heme cofactor that is essential for activity. First, supplementing cultures expressing NnlA with the heme precursor 5-ALA resulted in NnlA preparations that bound one *b*-heme per subunit (**Fig. 2, Table S2, and Fig. S2**). Reduction of the heme is required to initiate activity (**Fig. 4**). Finally, generation of the heme-depleted variant, NnlA H73A, results in loss of NNG degradation activity (**Fig. S5**).

A key goal of this study was to understand the need for an Fe^II^ source in the previous study. Without culture medium supplementation with 5-ALA, preparations of NnlA in the prior study had low heme occupancies (**Fig. 2**). While samples had enough heme-bound monomers to turnover NNG, the low occupancy precluded detection of the cofactor by UV-visible absorption spectroscopy. With the observation that Fe(NH_4_)_2_(SO_4_)_2_ reduced the NnlA heme (**Fig. S3**), it is most likely that this Fe^II^ source reduced a small quantity of heme-bound NnlA, all of which was responsible for the observed activity. We conclude that initiation of NNG degradation by NnlA is dependent on reduction of the heme and does not require the presence of non-heme Fe^II^.

Heme cofactors often mediate redox-dependent chemistries. For example, reductive degradation of the cyclic nitramine RDX is mediated by the heme cofactor of XplA (15–18). Furthermore, heme cofactors are well-established to activate O_2_,(46) presenting the possibility of a high-valent iron-oxo species participating in NNG decomposition. However, our experiments showed that activity does not require O_2_, precluding the possibility that O_2_ activation is required for NNG degradation (**Fig. 4**). In fact, O_2_ inhibited the NnlA-catalyzed decomposition of NNG (**Fig. 3**). Furthermore, Fe^II^-NnlA can perform at least 100 turnovers (**Table 2**), indicating that NNG is not reduced during degradation. Finally, complete determination of the nitrogen mass balance verified NH_4_^+^ as the second nitrogenous product of NNG decomposition by NnlA (**Table 2**). The combined results are consistent with NNG degradation by NnlA being a redox-neutral process.

The finding of an essential heme for NnlA activity now requires further understanding of its specific role. As discussed above, NnlA is predicted to be structurally homologous with PAS domains. In addition, we have established that NnlA required a heme cofactor for activity. We characterized NnlA as a dimer (**Fig. S1B**), and PAS domain proteins are often dimers or tetramers in solution (44, 61). In addition, suggested templates for structural homology modeling by SWISS-MODEL were nearly all structures of PAS domain proteins. Finally, the NnlA heme is capable of binding CO and NO to form stable complexes (**Fig. 3**). The NnlA heme does not stably bind O_2_ but is instead oxidized by O_2_. However, the structural homology model predicts that a critical distal tryptophan residue that stabilizes O_2_ binding in *Pa* Aer2 (60) is absent in NnlA and is replaced with a glutamine residue (**Fig. 6**). This may account for the differing reactivity of the NnlA heme compared to those of previously characterized heme-binding PAS domains that are O_2_ sensors. Regardless, the accumulated evidence supports assignment of NnlA as a PAS domain protein.

Typically, PAS domains are redox or O_2_ sensor domains (43, 44, 53). Activation of the domains by change in oxidation state or binding of O_2_ triggers downstream processes, such as kinase activity or chemotaxis. Therefore, one possibility is that the NnlA heme is a redox sensor that triggers organization of an allosteric active site for NNG degradation in NnlA (**Scheme 2**). Most PAS domain-containing subunits do not also contain the catalytic site. However, our results indicate that the NnlA monomer contains both the regulatory PAS domain as well as a catalytic domain. Alternatively, the active site may reside at the subunit interface thereby requiring oligomerization for activity. By this hypothesis, NnlA activity would be regulated by the redox environment, either by the presence of O_2_ or NO_2_^−^. As shown in **Fig. 3**, O_2_ oxidizes Fe^II^-NnlA, which inactivates the enzyme. This observation is consistent with the observation of less NNG degradation in oxic samples (**Fig. 4**). The oxidation of NNG by NO_2_^−^ (**Fig. 5**) indicates that NO_2_^−^ could also act in a regulatory fashion as a product inhibitor.

A second hypothesis is that the Nnla heme is repurposed as an active site instead of a regulatory site and therefore, the heme directly participates in NNG degradation. It was previously proposed that a general base deprotonates the NNG α-carbon, thereby, promoting β-elimination of NO_2_^−^ from NNG to form an imine (42). Subsequent hydrolysis of the imine intermediate results in formation of the degradation products NH_4_^+^ and glyoxylate. By our second hypothesis, the Fe^II^-heme could act as a Lewis acid to activate the nitro group and promote elimination of the NO_2_^−^ (**Scheme 2**). Differentiating between these two hypotheses will be required to identify the active site and enable future engineering efforts.

While the physiological function and biosynthetic pathway of NNG is still unknown, the finding of an NNG-degrading enzyme suggests that it is present in some abundance in the environment. It was previously shown that NNG is a potent inhibitor of the metabolic enzyme succinate dehydrogenase.(41) Additionally, NNG is isolectronic with 3-nitropropionate, a potent toxin produced by some fungi and plants.(62) It is reasonable to posit that NNG acts in some antibiotic function and that NnlA is a nitramine detoxification enzyme. However, the observation that NNG degradation by NnlA is product inhibited (or by the redox environment) would be unusual for a detoxification enzyme. Further studies of the role of this molecule and the ecological function of NnlA are warranted. Future studies may engineer NnlA to degrade nitramines of environmental concern, leveraging the mechanistic insight from this work.

## Acknowledgements

This research was made possible by funding from the United States Army Research Office under grant #W911NF2010286. We thank the UCF McNair Scholars Program for supporting the training and research of A.T. This work was supported in part by the Strategic Environmental Research and Development Program (SERDP) under project WP20-1151. Oak Ridge National Laboratory is managed by UT-Battelle, LLC, for the U.S. Department of Energy under contract no. DE-AC05-00OR22725. We also thank Dr. Matthew Rex and Bhavini Goswami for their assistance in LC–MS.

## Schemes

**Scheme 1.**
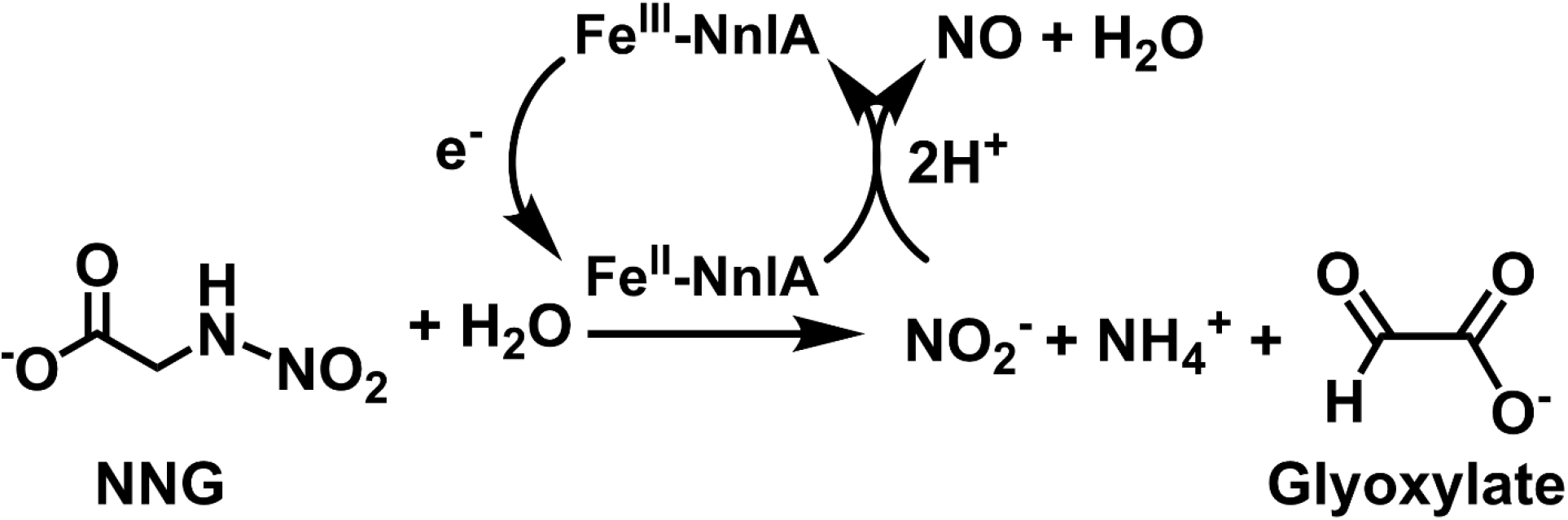
Summary of NnlA activity.

**Scheme 2.**
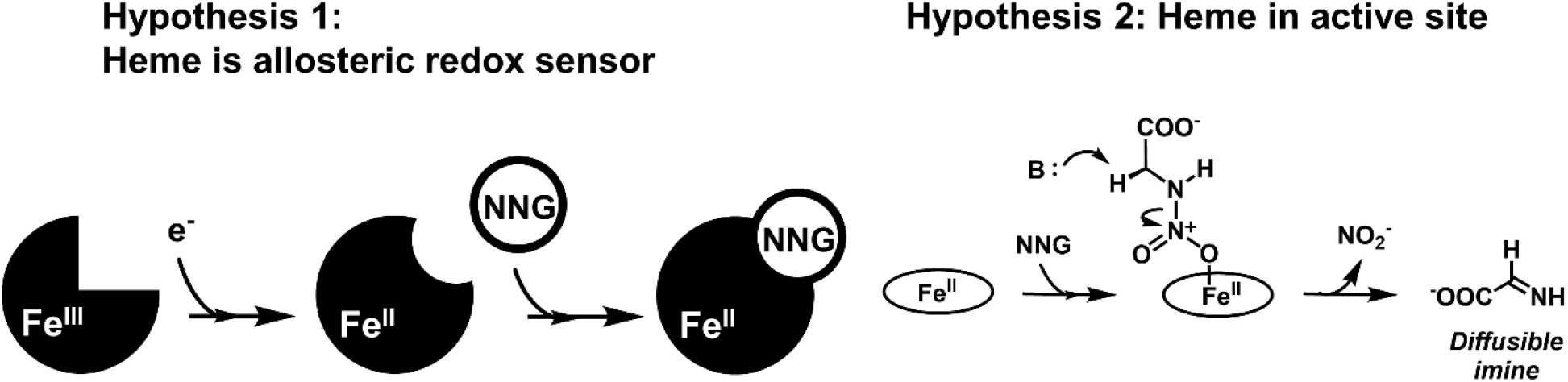
Proposed roles of heme cofactor in NnlA.

