## Supplemental_Figures_and_Tables for "Reduction of a heme cofactor initiates N-nitroglycine degradation by NnlA"

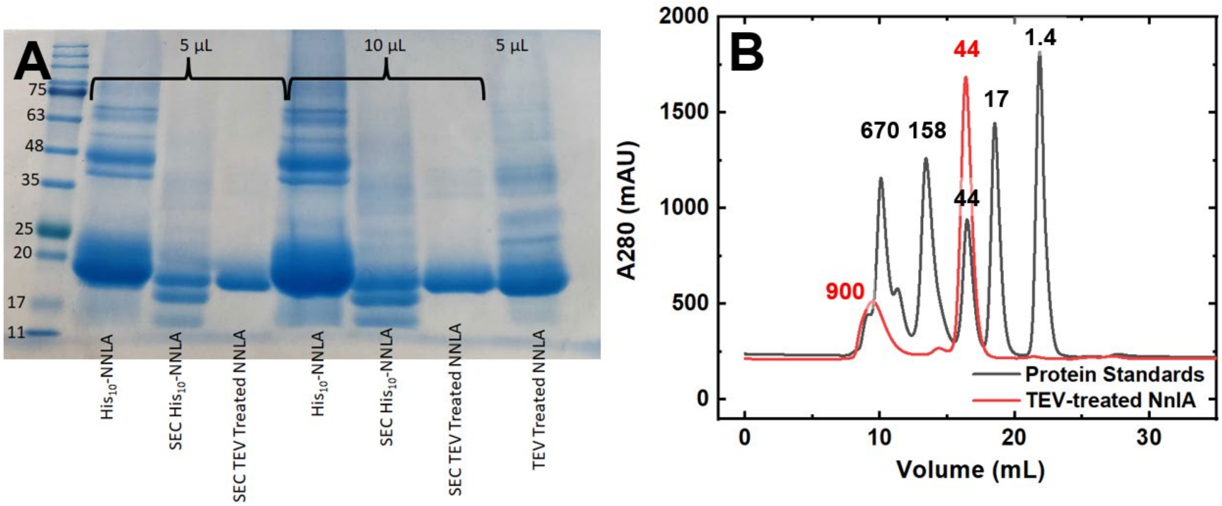


**Fig. S1.** NnlA protein size determination as determined by A) SDS-PAGE of purified NnlA fractions or B) analytical size exclusion chromatography of purified $His_{10}$-NnlA (red trace). Black trace in panel B shown protein molecular weight standards.

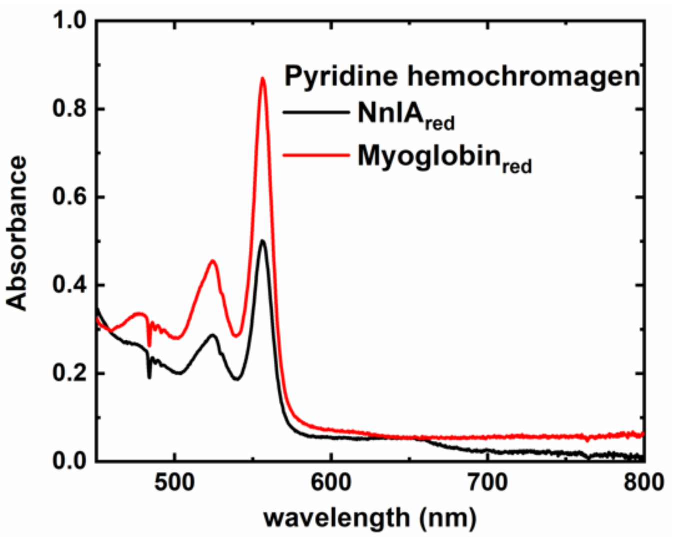


**Fig. S2**. UV-visible absorption spectra of the pyridine hemochromogen assay with reduced NnlA (red trace) and myoglobin (black trace). Final concentrations were 33 µM myoglobin and 23 µM NnlA in 0.11 M NaOH and pyridine with sodium dithionite.

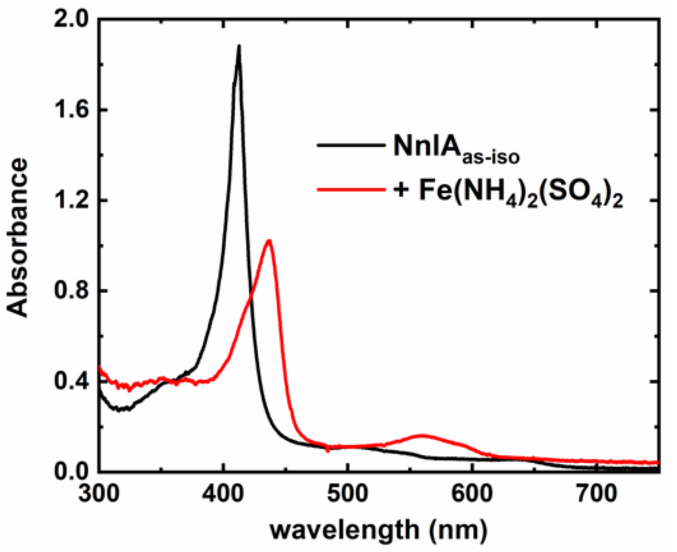


**Fig. S3.** UV-visible absorption spectra illustrating the reduction of NnlA$_{as\text{-}iso}$ with $Fe(NH_4)_2(SO_4)_2$. Treatment of 5.5 µM NnlA$_{as\text{-}iso}$ (black trace) with 26 µM $Fe(NH_4)_2(SO_4)_2$ and incubated at room temperature (red trace).

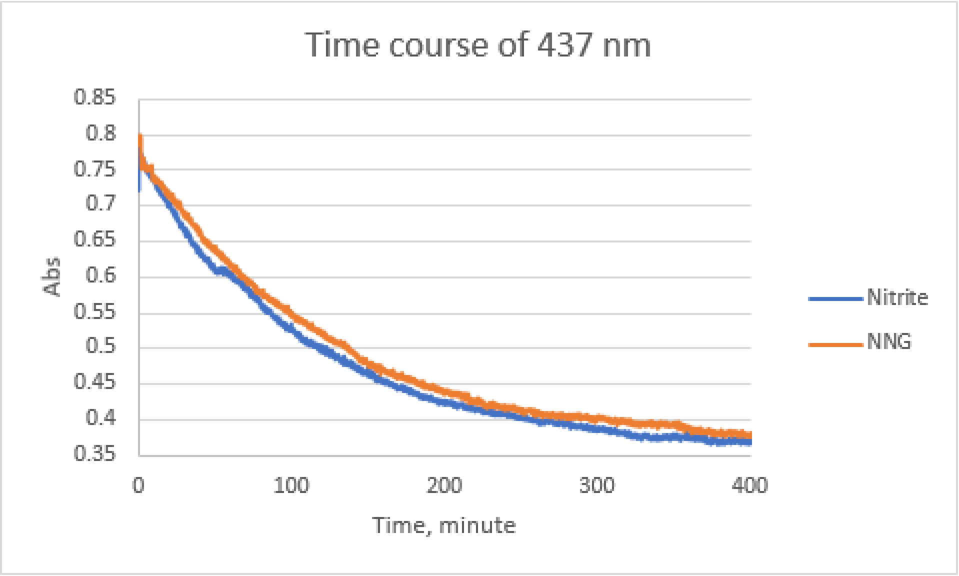


**Fig. S4**. Time course for oxidation of $Fe^{II}$-NnlA by $NO_2^-$ and NNG as monitored by single wavelength UV-visible absorption spectrometry.

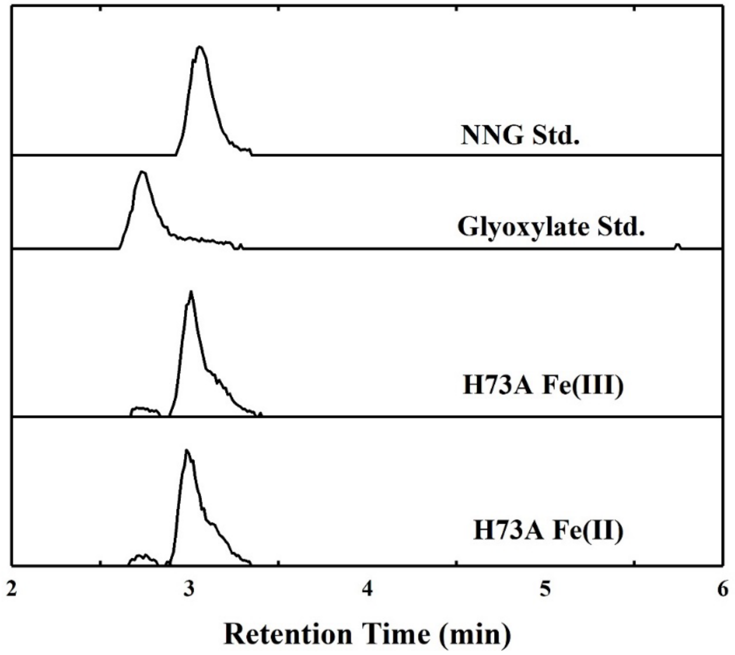


**Fig. S5.** LC-MS analysis of H73A activity: H73A control (10 µM H73A NnlA, + 400 µM NNG) vs reduced H73A Fe (II) (10 µM Sodium Dithionite + 10 µM H73A NnlA, + 500 µM NNG), both samples were prepared in nitrogen atmosphere glovebox, incubated for 30 minutes at 20°C in 10 mM tricine buffer, pH 8.

| **Table S1. H73A mutant primers** | | |
|---|---|---|
| | Forward primer | Reverse Primer |
| H73A | GCCCGTCGTGCAATCACGAAT | GCGATTATGAATTTCATCCCCAAT |

| Table S2. Iron analysis of wild-type, TEV-cleaved wild-type, and H73A NnlA.[a] | | | | |
|---|---|---|---|---|
| **Sample** | **[Non-heme iron](μM)[b]** | **[iron]total (μM)[c]** | **[NnlA monomer] (μM)[d]** | **[iron]total/[NnlA monomer](μM)** |
| **Wild-type NnlA** | ND[d] | 260 ± 30 | 300 ± 60 | 0.87 ± 0.21 |
| **TEV cleaved Wild-type NnlA** | 2.5 | 28.2 ± 0.4 | 30 ± 1 | 0.95±0.04 |
| **H73A NnlA** | 91 ± 18 | 49 ± 11 | 140 ± 1 | 0.35 ± 0.08 |

[a]Values reported as averages and standard deviations of 3 trials.
[a]Quantified by ferrozine assay.
[b]Quantified using a modified assay reported by Fish.
[c]Quantified by BCA assay.
[d]Not detected

| Table S3. Total dissolved metal concentrations of Ni-HTC purified $His_{10}$-NnlA samples by ICP-MS.[a] | |
|---|---|
| **Analyte** | **[Metal]/[NnlA monomer]** |
| **Fe** | 0.44 |
| **Ni** | 1.86 |
| **Co** | 0.0009 |
| **Cu** | 0.17 |
| **Zn** | 0.15 |
| **Mo** | 0.0005 |
| **Mn** | 0 |

[a]Samples were prepared by treating protein samples with trichloroacetic acid and 2% nitric acid and pelleting the denatured protein prior to analysis.

**Table S4.** $NO_2^-$ and $NO_3^-$ concentrations of samples by ion chromatography or Griess assay.[a]

| **Sample** | **$[NO_2^-]_{IC}$ (μM)[b]** | **$[NO_2^-]_{Griess}$ (μM)[c]** | **$[NO_3^-]_{IC}$ (μM)[b]** |
|---|---|---|---|
| **$Na_2S_2O_4$ + NNG[a]** | 28 ± 10 | 25 ± 11 | 6.3 ± 0.6 |
| **$Fe^{II}$-NnlA + NNG[a]** | 136 ± 12 | 165 ± 13 | 6.9 ± 0.6 |

[a]Samples contained 250 μM NNG in 10 mM tricine buffer at pH 8.0 either with or without 5 μM $Fe^{II}$-NnlA and incubated for 30 minutes at room temperature under anaerobic conditions.
[b]Detected by ion chromatography
[c]Detected by Griess assay
